# Postpartum corticosterone and fluoxetine shift the tryptophan-kynurenine pathway in dams

**DOI:** 10.1101/2021.02.11.430473

**Authors:** Wansu Qiu, Kimberly A. Go, Yvonne Lamers, Liisa A. M. Galea

**Author notes:** Corresponding Author: 2215 Wesbrook Mall, Vancouver, BC, Canada, V6T 1Z3.

## Abstract

Perinatal depression (PND) affects 15% of mothers. Selective serotonin reuptake inhibitors (SSRIs) are currently the first-line of treatment for PND but are not always efficacious. Previously, we found significant reductions in plasma tryptophan concentrations and higher hippocampal proinflammatory cytokine, IL-1b levels, due to maternal SSRI treatment. Both inflammation and tryptophan-kynurenine metabolic pathway (TKP) are associated with SSRI efficacy in individuals with major depressive disorder (MDD). TKP is divided into neuroprotective and neurotoxic pathways. Higher metabolite concentrations of the neurotoxic pathway are associated with depression onset and implicated in SSRI efficacy. Metabolites in TKP were investigated in a rodent model of de novo postpartum depression (PPD) given treatment with the SSRI, fluoxetine (FLX). Dams were administered corticosterone (CORT) (40mg/kg, s.c.), and treated with the SSRI, fluoxetine (FLX) (10mg/kg, s.c.), during the postpartum for 22 days after parturition. Plasma TKP metabolite concentrations were quantified on the last day of treatment. Maternal postpartum CORT increased neurotoxic metabolites and co-enzyme/cofactors in dams (3-hydroxykynurenine, 3-hydroxyanthranilic acid, vitamin B2, flavin adenine dinucleotide). The combination of both CORT and FLX shifted the neuroprotective-to-neurotoxic ratio towards neurotoxicity. Postpartum FLX decreased plasma xanthurenic acid concentrations. Together, our data indicate higher neurotoxic TKP expression due to maternal postpartum CORT treatment, similar to clinical presentation of MDD. Moreover, maternal FLX treatment showed limited efficacy to influence TKP metabolites, which may correspond to its limited efficacy to treat depressive-like endophenotypes. Overall suggesting changes in TKP may be used as a biomarker of de novo PPD and antidepressant efficacy and targeting this pathway may serve as a potential therapeutic target.

**Highlights:**

- Tryptophan-kynurenine pathway (TKP) is altered by postpartum corticosterone (CORT)
- Postpartum CORT upregulated neurotoxic more metabolites (3HK, 3HAA)
- Postpartum fluoxetine (FLX) increased xanthurenic acid concentrations
- Postpartum CORT and FLX together shifted the TKP balance towards neurotoxicity

## 1. Introduction

Perinatal depression (PND) is a heterogeneous disease, in which symptoms and remission vary depending on depression onset timing and depression history (reviewed in Qiu et al., 2020b). The first three months postpartum is a period of high susceptibility for first time psychiatric disease in females but not in males (Munk-Olsen et al., 2006). Postpartum depression (PPD) affects 15% of females with approximately 40% of these cases being individuals experiencing depressive symptoms for the first time (Wisner et al., 2013). Maternal postpartum corticosterone (CORT) is used to model *de novo* PPD in rodents (reviewed in Qiu et al., 2020b). This model mimics elevated cortisol levels seen in those women with depression symptoms postpartum (Iliadis et al., 2015). Maternal postpartum CORT consistently induces various depressive-like endophenotypes, such as reduced maternal care behaviour, increased passive-coping behaviour, and reduced hippocampal integrity (Workman et al., 2016; Gobinath et al., 2018; Qiu et al., 2020a).

Inflammation and altered metabolism in the tryptophan-kynurenine pathway (TKP) are associated with major depressive disorder (MDD) and PND (Corwin et al., 2008; Maes et al., 2011; Haapakoski et al., 2015). The immune system and TKP are related as proinflammatory cytokines, such as interferon (IFN)-γ and interleukin (IL.)-I β, can increase enzymatic conversion of tryptophan into its downstream metabolite, kynurenine (reviewed by Maes et al., 2011). Higher expression of proinflammatory cytokines such as IL-6, tumor necrosis factor (TNF)-α, IFN-γ, and IL-1β are associated with both MDD and PND (Corwin et al., 2008; Haapakoski et al., 2015). TKP can be divided into either a “neurotoxic” pathway or a “neuroprotective” pathway and higher activation in the neurotoxic branch is seen in MDD and PND (see Figure 1A; reviewed in Maes et al., 2011). Curiously, PND is associated with higher kynurenic acid (KYNA) (Teshigawara et al., 2019), but MDD is associated with reduced KYNA (reviewed by Maes et al., 2011). Previously, we found reduced cytokine (TNF-α and IFN-γ) expression and elevated concentrations of a TKP enzyme cofactor (vitamin B6) due to postpartum CORT (Qiu et al., 2020a), indicating an altered immune system and TKP in this model of *de novo* PPD.

**Figure 1.**
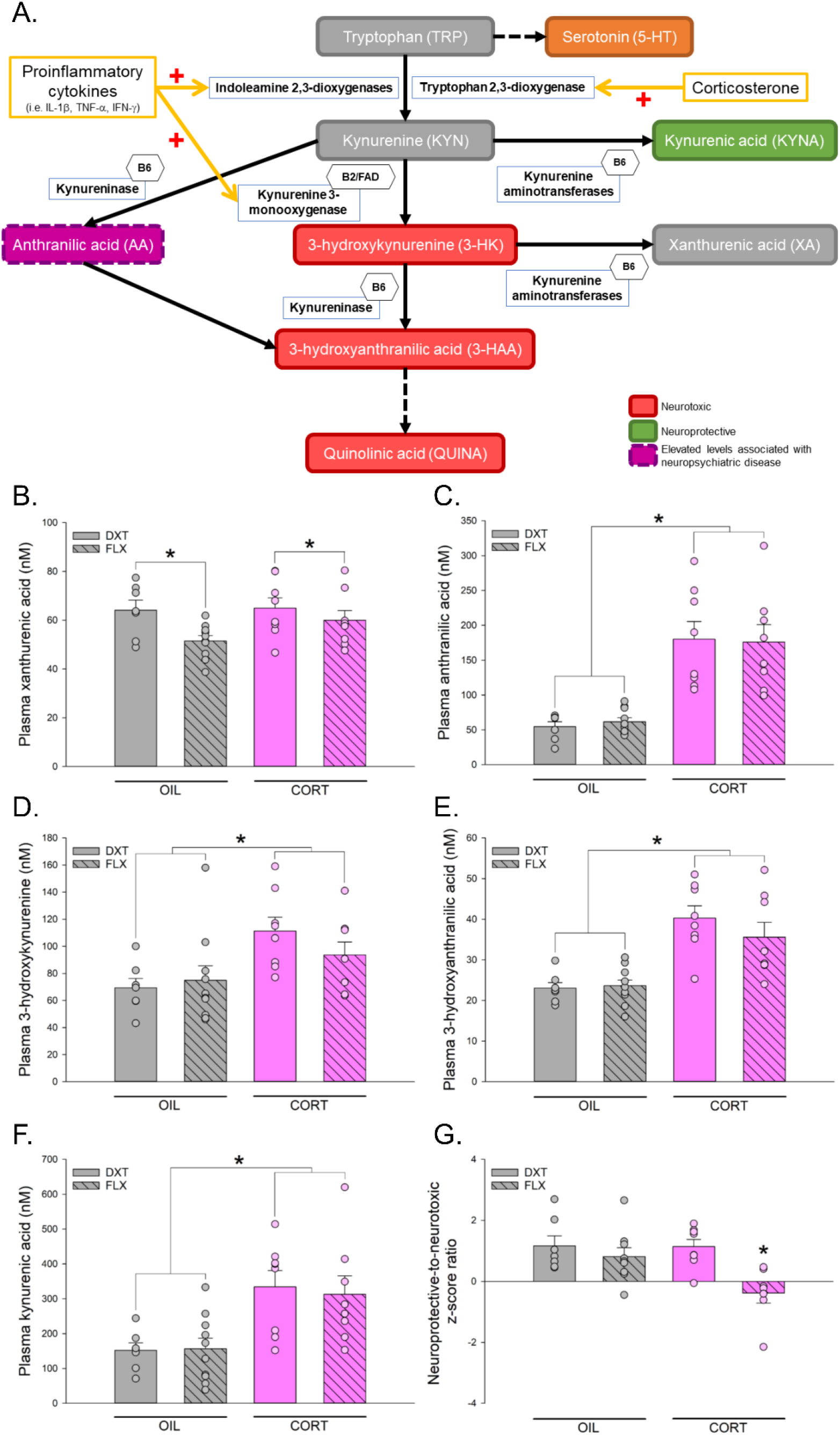
Effects of maternal corticosterone (CORT) and fluoxetine (FLX) on plasma tryptophan-kynurenine pathway metabolites. **A.** Scheme of the tryptophan-kynurenine pathway (TKP). **B.** FLX decreased plasma xanthurenic acid concentrations. **C.-F.** CORT treatment significanly increased plasma anthranilic acid concentrations **(C.)**, plasma 3-hydroxykynurenine concentrations **(D.)**, plasma 3-hydroxyanthranilic acid concentrations **(E.)**, and plasma kynurenic acid concentrations **(F.)**. **G.** CORT+FLX significantly decreased the neuroprotive-to-neurotoxic z score ratio compared to all other groups. \**p* < 0.05. Data represented in means + standard error of the mean, overlaid with individual data points, n’s = 7-10 per group.

Selective serotonin reuptake inhibitors (SSRIs) are the first line treatment for PND although efficacy of SSRIs varies with timing of depression onset as those with postpartum onset have lower efficacy than those with antenatal onset (Cox et al.,2016). Both inflammation and TKP are implicated in antidepressant efficacy both in humans (Syed et al., 2018; Sun et al., 2020) and in rodents (Qiu et al., 2020a). Individuals with MDD who are non-responsive to SSRI treatment show either no change or an elevation of circulating cytokine levels (Syed et al., 2018) and display higher baseline tryptophan metabolism defined by kynurenine to tryptophan ratio (Sun et al., 2020). Previously, limited efficacy of the SSRI, fluoxetine (FLX), to decrease depressive-like behaviour, was commensurate with increased levels of hippocampal IL-1β and decreased circulating tryptophan concentrations (Qiu et al., 2020a). Thus, tryptophan catabolism via TKP may be disrupted by CORT and could be a potential biomarker of SSRI efficacy, and the present study sought to understand this relationship. Here, using a model of *de novo* PPD, we investigated the effects of maternal postpartum CORT, to induce depressive-like endophenotypes, and concurrent FLX to understand how these treatments may affect metabolites in the TKP. We hypothesized that maternal postpartum CORT treatment will alter TKP metabolite concentrations and that postpartum FLX will have limited efficacy to inhibit these effects.

## 2. Methods

Forty adult female Sprague-Dawley rats and six male rats (2.5 months old) were purchased from Charles River (Montreal, QC, Canada). Males were used as breeders, and co-housed with two females until pregnancy (determined via sperm in daily vaginal lavage samples). All pregnant rats were undisturbed until parturition. Thirty-seven females were included in the study. Protocols were in accordance with ethical guidelines set by the Canadian Council for Animal Care and were approved by the University of British Columbia Animal Care Committee.

All methods are previously described in Qiu et al. (2020a). Briefly, all dams received daily injections postpartum, starting from two days after birth, for 22 days. All injections were given subcutaneously (s.c.) and occurred between 08:00-10:00. Dams received CORT (40mg/kg, s.c.; Sigma-Aldrich, St. Louis, MO, USA) or its vehicle (OIL; sesame oil + 10% EtOH, 1ml/kg, s.c.) and either FLX (10mg/kg, s.c.; Sequoia Research Products, Pangbourne, UK) or its vehicle, dextrose (DXT; 5% dextrose in sterile water, 1ml/kg, s.c.). Dams were randomly assigned to one out of four treatment groups (OIL+DXT, n = 9; OIL+FLX, n = 10; CORT+DXT, n = 9; and CORT+FLX, n = 9). All animals underwent maternal observations from postnatal day (PD) 2-8 and then testing in the forced swim test on PD21-22 and these data were published previously (Qiu et al., 2020a). We found postpartum CORT reduced total nursing and increased time off the nest and postpartum FLX reversed these effects in the early postpartum. However, on PD23, we found in these rats that postpartum CORT increased immobility in the FST that was not rescued by FLX, an effect that was consistent with past research (Workman et al., 2016; Gobinath et al., 2018). At least 2 hours after the last injection and immediately after the FST, all animals were euthanized by rapid decapitation, brains were extracted and trunk blood was collected within three minutes of touching the cage. Plasma was collected in cold EDTA (made in house) coated microcentrifuge tubes, immediately placed on ice to help preserve contents in plasma, and centrifuged 4h later for 10min at 4°C. Data for 5 samples were not collected due to human error. We also examined inflammation in the periphery, hippocampus and frontal cortex and these results have been published elsewhere (Qiu et al., 2020a). Briefly, we found that postpartum FLX elevated hippocampal IL-1β and circulating CXCL1 and postpartum CORT reduced IFN-γ and TNF-α. Here we examined TKP metabolites.

We quantified kynurenine (KYN), kynurenic acid (KYNA), xanthurenic acid (XA), anthranilic acid (AA), 3-hydroxykynurenine (3-HK), 3-hydroxyanthranilic acid (3-HAA), riboflavin (vitamin B2) and its cofactor, flavin adenine dinucleotide (FAD), using isotope dilution liquid chromatography coupled with tandem mass spectrometry (ABSciex API4000; AB SCIEX, Framingham, MA, U.S.A.) based on a modified method by Midttun et al. (2005). The intra- and inter-assay CVs for the quantitation of these analytes were all < 7%. Using a novel exploratory approach, a neuroprotective-to-neurotoxic z score ratio was calculated to account for the balance between multiple neuroprotective metabolites or neurotoxic metabolites/cofactors: (zKYNA + zAA) / (z3-HK + z3-HAA + zB2). All metabolites and metabolite ratio data were analyzed using two-way general linear model of analysis of covariance (ANCOVA) with CORT (CORT or OIL) and FLX (FLX or DXT) as between-subject factors and estrous cycle stage as covariate, unless otherwise specified. Effect sizes were reported as □_p_^2^ or Cohen’s *d*. Post hoc comparisons used Newman-Keuls. *A priori* comparisons were subjected to Bonferroni correction. Pearson’s product-moment correlations were conducted between behavior, and cytokine data, adrenal mass, and TKP metabolites. All statistical analyses were analyzed using Statistica software (v. 9, StatSoft, Inc., Tulsa, OK, USA). The covariate of estrous cycle stage and body mass did not significantly affect any analyses (all *p*’s ≥ 0.097). Outliers were removed when two standard deviations away from the mean, and this happened twice in the neuroprotective-to-neurotoxic ratio. Assumptions of normal distributions and homogeneity of variance were meet and verified with tests for normality and Barlett’s test for all dependent variables.

## 3. Results

Maternal FLX treatment decreased XA compared to vehicle DXT treatment (main effect of FLX: F(1, 29) = 6.009, *p* = 0.020, □_p_^2^ = 0.172; Figure 1B), with no other significant effects (all *p*’s ≥ 0.296).

Maternal CORT increased AA (main effect of CORT: F(1, 29) = 44.506, *p* < 0.001, □_p_^2^ = 0.605), 3-HK (F(1, 29) = 9.282,*p* = 0.005, □_p_^2^ = 0.242), and 3-HAA (F(1, 29) = 33.693,*p* < 0.001, □_p_^2^ = 0.537) concentrations compared to vehicle-treated animals (Figure 1C-E), with no other significant effects (all *p*’s ≥ 0.253). Maternal CORT also increased plasma KYNA (F(1, 29) = 17.731, *p* < 0.001; □_p_^2^ = 0.379; Figure 1F) with no other significant effects (all *p*’s ≥ 0.747). CORT+FLX significantly lowered the neuroprotective-to-neurotoxic z score ratio compared to all other treatment groups, indicating a shift towards the neurotoxic pathway *(a priori* all *p*’s ≤ 0.007; Cohen’s *d* = 1.785 compared to OIL+DXT, Cohen’s *d* = 1.372 compared to OIL+FLX, Cohen’s *d* = 1.963 compared to CORT+DXT; interaction between CORT and FLX, F(1, 27) = 3.906, *p* = 0.058, □_p_^2^ = 0.126; main effect of CORT, F(1, 27) = 4.265, *p* = 0.049, □_p_^2^ = 0.136; main effect of FLX, F(1, 27) = 10.055, *p* = 0.004, □_p_^2^ = 0.271, Figure 1G). There were no other significant effects on KYN or other metabolite concentrations (*p* = 0.410)

Vitamin B2 and its cofactor, FAD, are critical for the enzymatic conversion of KYN to the neurotoxic metabolite 3-HK, playing an important role in the neurotoxic pathway of TKP. Maternal CORT increased plasma vitamin B2 (main effect of CORT: F(1, 29) = 5.070, *p* =

0.032, □_p_^2^ = 0.149), with no other significant effects (all *p*’s ≥ 0.254, Figure 2A). CORT+FLX had significantly higher FAD than all other groups (all *p*’s ≤ 0.021; Cohen’s *d* = 1.718 compared to OIL+DXT, Cohen’s *d* = 2.166 compared to OIL+FLX, Cohen’s *d* = 1.325 compared to CORT+DXT; interaction between CORT and FLX, F(1, 29) = 3.888, *p* = 0.050, □_p_^2^ = 0.118; main effect of CORT, F(1, 29) = 13.432, *p* = 0.001, □_p_^2^ = 0.316; no significant main effect of FLX, *p* = 0.151, Figure 2B).

**Figure 2.**
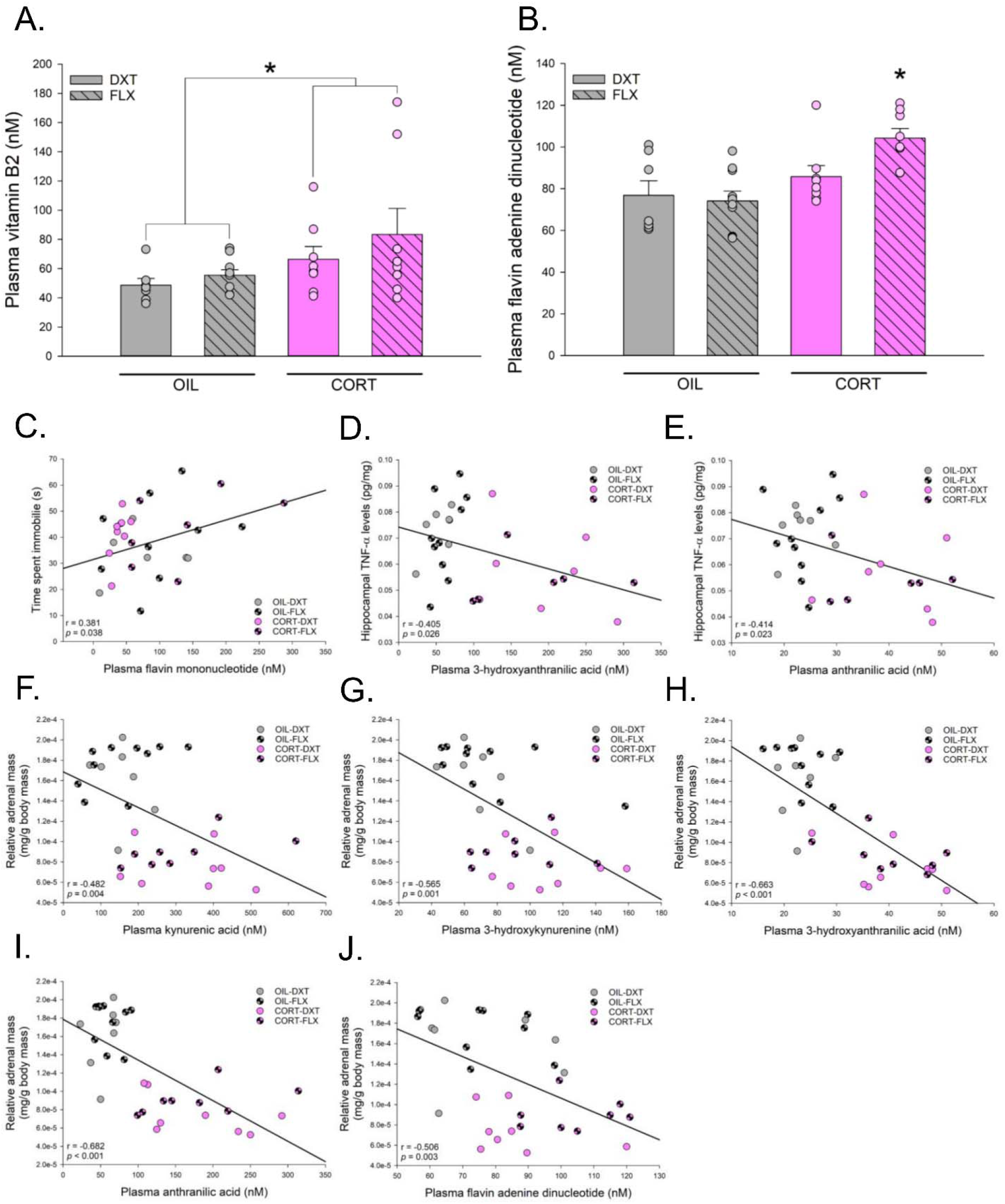
Effects of maternal corticosterone (CORT) and fluoxetine (FLX) treatment on tryptophan-kynurenine pathway coenzyme and cofactor. **A.** CORT significantly increased plasma vitamin B2 concentrations. **B.** CORT+FLX significantly increased plasma flavin adenine dinucleotide (FAD) concentrations compared to all other groups. **C.** Plasma flavin mononucleotide positively correlated time spent immobile in the forced swim test. **D.** Plasma 3-hydroxyanthranilic acid (3-HAA) negatively correlated with hippocampal tumor-necrosis factor (TNF)-α levels. **E.** Plasma anthranilic acid (AA) negatively correlated with hippocampal TNF-α levels. **F.** Plasma kynurenic acid (KYNA) negatively correlated with relative adrenal mass. **G.** Plasma 3-hydroxykynurenine (3-HK) negatively correlated with relative adrenal mass. **H.** Plasma 3-HAA negatively correlated with relative adrenal mass. **I.** Plasma AA negatively correlated with relative adrenal mass. **J.** Plasma flavin adenine dinucleotide (FAD) negatively correlated with relative adrenal mass. \**p* < 0.05. Data represented in means + standard error of the mean, overlaid with individual data points, n’s = 7-10 per group.

Plasma flavin mononucleotide positively correlated with time spent immobile in the forced swim test (r = 0.381, *p* = 0.038) (Figure 2C). Hippocampal levels of the proinflammatory cytokine, TNF-α negatively correlated with both 3-HAA (r = −0.405, *p* = 0.026) and AA (r = −0.414, *p* = 0.023), Figure 2D-E. Relative adrenal mass negatively correlated with KYNA, (r = −0.482, *p* = 0.004), 3-HK (r = −0.565, *p* = 0.001), 3-HAA (r = −0.663, *p* = < 0.001), AA (r = −0.682, *p* < 0.001), and FAD (r = −0.506,*p* = 0.003). There were no other significant correlations (all *p*’s ≥ 0.064).

## 4. Discussion

Maternal postpartum CORT increased two neurotoxic TKP metabolites, 3-HK and 3-HAA, and one coenzyme of the neurotoxic pathway, vitamin B2. However, postpartum CORT also increased two neuroprotective metabolites, KYNA and AA, without significantly affecting KYN concentrations. Further, maternal postpartum CORT with postpartum FLX shifted the neuroprotective-to-neurotoxic balance towards neurotoxicity and increased FAD, a coenzyme to drive conversion to the neurotoxic pathway. Maternal postpartum FLX treatment alone decreased plasma XA concentration. Together, these data indicate maternal postpartum CORT upregulated TKP towards the neurotoxic pathway, and this effect together with FLX increased neurotoxic signalling in the TKP.

Maternal postpartum CORT increased plasma concentrations of neurotoxic metabolites, 3-HK and 3-HAA, which is consistent with findings using inflammation-induced depression models (Parrott et al., 2016; Tao et al., 2020). Inflammation-induced depression, via lipopolysaccharide, results in higher concentrations of 3-HK in the frontal cortex and hippocampus of male mice (Tao et al., 2020). Lipopolysaccharide-induced depressive endophenotypes are dependent on the activation of the neurotoxic pathway of TKP, via 3-HK, at least in male mice (Parrot et al., 2016). Maternal CORT also increased the neuroprotective metabolite AA, which is consistent with findings showing elevated peripheral concentrations in both male and female individuals with high risk for MDD and in an animal model of depression (Sakurai et al., 2020). Together these results suggest that maternal postpartum CORT activates the neurotoxic pathway of TKP, which shares similarities to other models of depression as well as in individuals with MDD, and therefore may prove to be an effective therapeutic target in the future.

KYNA concentrations are reduced in MDD (reviewed by Maes et al., 2011), however a meta-analysis revealed that lower KYNA concentrations with MDD were found when there was a higher percentage of male patients (Ogyu et al., 2018), indicating a possible sex difference in this effect. Indeed, higher gestational KYNA concentrations were found in women with the onset of postpartum depressive symptoms (Teshigawara et al., 2019). This latter finding is consistent with our findings that maternal postpartum CORT increased depressive-like endophenotypes (Qiu et al., 2020a) and in the present study increased plasma KYNA, a neuroprotective metabolite. It is possible the increases in both neuroprotective and neurotoxic metabolites by maternal postpartum CORT reflects higher TKP activation, which is upregulated by glucocorticoids in both rodents and humans (reviewed in Maes et al., 2011). Specifically, CORT can upregulate tryptophan 2,3-dioxygenase activity, increasing the breakdown of tryptophan to kynurenine (reviewed by Maes et al., 2011). This effect occurs as a first step in the TKP, prior to the division of metabolites into either the neuroprotective or neurotoxic pathway. Regardless, postpartum CORT increased activation of the neurotoxic pathway via elevated concentrations of two neurotoxic metabolites (3-HK and 3-HAA), and the neurotoxic branch coenzyme vitamin B2, and its cofactor FAD. Thus, higher activation of the neurotoxic branch may have contributed towards the CORT-induced depressive-like endophenotypes reported previously in these same animals (Qiu et al., 2020a). In this regard it is notable that FAD was positively correlated with passive coping in the FST. Future studies could use this as therapeutic target and/or increase the efficacy of FLX by targeting this pathway.

Previously, we reported in these same animals, a significant decrease in plasma tryptophan concentration and higher levels of the proinflammatory cytokine, IL-1β, in the maternal hippocampus as well as higher peripheral CXCL1 levels due to FLX treatment in the same animals (Qiu et al., 2020a). Higher levels of IL-1β can also upregulate TKP (reviewed in Maes et al., 2011), so this may have contributed to the current findings. Maternal postpartum FLX decreased plasma XA concentration, which is consistent with data in individuals with MDD (Colle et al., 2020). XA has not been specifically described as either neuroprotective or neurotoxic, but its downstream effects on the glutaminergic system have been implicated in psychiatric diseases (reviewed Fazio et al., 2018). Moreover, maternal postpartum FLX with CORT increased FAD, a cofactor in the neurotoxic branch, and shifted the balance of the neuroprotective-to-neurotoxic ratio towards the neurotoxic branch. Previously, in these same animals, postpartum FLX failed to reverse depressive-like endophenotypes (Qiu et al., 2020a). Further, we found significant negative correlations between hippocampal TNF-α levels and plasma 3-HAA and AA concentrations. Although this finding is partially inconsistent with a study in MDD individuals where TNF was positively associated with plasma TKP (Haroon et al., 2020), this other study used sex as covariate and did not consider treatment effects in their analyses. The present findings coupled with the results of higher IL-1β and reduced TRP concentrations with maternal postpartum FLX (Qiu et al., 2020a) may have contributed towards the limited efficacy of FLX previously reported in this model of *de novo* PPD.

It is important to note that in this study we examined peripheral levels of TKP metabolites and it is very possible that brain levels of TKP metabolites may be different. However, Haroon and colleagues (2020) found that peripheral plasma TKP activation and metabolites were positively correlated with cerebrospinal fluid TKP activation and metabolites, indicating some consistency between peripheral and central levels. Using postpartum CORT, we see several depressive-like endophenotypes that overlap with the human presentation of *de novo* postpartum depression (reviewed in Qiu et al., 2020b). As stated earlier, the efficacy for antidepressants is lower for those with postpartum depression compared to antepartum (Cox et al., 2016). Further, women with PPD, show early remission with SSRI treatment, which fades with longer treatment (Sharp et al., 2010), suggesting short-term but not long-term efficacy of SSRIs. Our own findings mirror the human data as we have found that SSRIs reverse the maternal care deficits seen with postpartum CORT in the short term (PD 2-8) but do not reverse the effects on passive coping or on hippocampal integrity in the long term (PD 23) (Workman et al., 2016; Gobinath et al., 2018, and Qiu et al., 2020a; reviewed by Qiu et al., 2020b). As perinatal depression is most common in women with a previous history of depression it would be important to determine if other PND models of antenatal depression similar effects on the TKP.

## Conclusion

Overall, high maternal postpartum CORT, which models *de novo* depression onset during the postpartum, upregulated metabolites in the neurotoxic pathway of the TKP, consistent with the profiles of TKP in MDD. Maternal postpartum CORT also led to higher concentrations of neuroprotective metabolites, KYNA and AA, both of which have been associated with either MDD or PPD. This suggests that the association between TKP metabolites and depression may be an important therapeutic target. Maternal postpartum FLX commensurate with maternal CORT increased TKP activation by increasing FAD concentrations and shifting the neuroprotection-to-neurotoxic balance towards the neurotoxic pathway of TKP. These results indicate that higher activation of metabolites of the neurotoxic TKP pathway may be a biomarker of depression and targeting this pathway may lead to novel therapeutic targets for PND and MDD.

## Acknowledgements

The authors would like to thank all the wonderful staff members at the Centre for Disease Modelling animal facility at the University of British Columbia for the continuous support and help throughout the project. Financial support for this research was provided by an operating grant from the Canadian Institutes for Health Research (MOP142308) to LAMG. WQ is supported by a 4-year Fellowship granted by the University of British Columbia, Faculty of Medicine, Graduate Program in Neuroscience, and the University of British Columbia Institute of Mental Health Marshalls Scholars Program in Mental Health, Department of Psychiatry.

